# Intraspecific Competition and the Promotion of Ecological Specialization

**DOI:** 10.1101/2021.09.04.458988

**Authors:** Abdel H. Halloway, Joel S. Brown

**Affiliations:** University of Illinois at Chicago; Moffitt Cancer Center, University of Illinois at Chicago

## Abstract

The evolution of ecological specialization can be summed up in a single question: why would a species evolve a more-restricted niche space? Various hypotheses have been developed to explain the promotion or suppression of ecological specialization. One hypothesis, competitive diversification, states that increased intraspecific competition will cause a population to broaden its niche breadth. With individuals alike in resource use preference, more individuals reduce the availability of preferred resources and should grant higher fitness to those that use secondary resources. However, recent studies cast doubt on this hypothesis with increased intraspecific competition reducing niche breadth in some systems. We present a game-theoretic evolutionary model showing greater ecological specialization with intraspecific competition under specific conditions. This is in contrast to the competitive diversification hypothesis. Our analysis reveals that specialization can offer a competitive advantage. Largely, when facing weak competition, more specialized individuals are able to acquire more of the preferred resources without greatly sacrificing secondary resources and therefore gain higher fitness. Only when competition is too great for an individual to significantly affect resource use will intraspecific competition lead to an increased niche breadth. Other conditions, such as a low diversity of resources and a low penalty to specialization, help promote ecological specialization in the face of intraspecific competition. Through this work, we have been able to discover a previously unseen role that intraspecific competition plays in the evolution of ecological specialization.

## Introduction

Through the process of adaptation and speciation, evolution by natural selection has produced a multitude of species of varying forms, all presumably optimized to their environment (Darwin, 1859). Despite the diversity of resources and environments available to each organism, each species is restricted to a subset of them. This “place” to which a species belongs is known as the niche (Leibold, 1995). Grinell first defined the niche as the requirements necessary for an organism to survive; this was later described by Hutchinson as the *n*-dimensional space within a system of *n*-dimensional ecological axes in which a species’ population can persist, its fundamental niche (hereafter, usage of the word niche refers to the fundamental niche unless otherwise specified) (Grinell, 1917; Hutchinson, 1957). This fundamental niche is intrinsic to the organism, often resistant to eco-evolutionary changes (Holt and Gaines, 1992; Wiens et al., 2010). Not only does niche space among species vary in position along the ecological axes but also in shape and size. Species which are said to have a smaller niche space are more restricted ecologically and said to be specialized compared to those with larger spaces. The existence of species with smaller niche space seems to be a paradox. Less specialized species with greater niche space should have higher fitness as they have access to more resources and are less vulnerable to extinction (REFs). And yet, specialization exists. This raises the question: why would a species evolve to restrict the environments in which it can live or the resources it can consume?

Many theories have been brought up as to why a species may specialize (Futuyma and Moreno, 1988). These include environmental constancy, trade-offs (Kotler and Mitchell, 1995; McNickle et al., 2016), genetic and phenotypic constraints (Futuyma et al., 1993; Futuyma et al., 1995; DeWitt et al., 1998), adaptation in and to a heterogeneous landscape (Holt and Gaines, 1992), co-evolution among mutualists (Fleming and Holland, 1998; Bronstein, 2009), and predation (Jeffries and Lawton, 1984). One factor given considerable attention is competition (Diamond, 1978). In particular, interspecific competition between species is assumed to promote specialization. Essentially, the presence of other species removes potential niche space from a focal species, which goes on to specialize on the remaining niche space and eventually has its previously realized niche become its fundamental niche (Van Valen, 1965; Cox and Ricklefs, 1977; Bolnick et al., 2010). This has been borne out in both theoretical (MacArthur and Levins, 1964; Slatkin, 1980) and experimental studies (reviewed in Araújo et al., 2011). Under this perspective, the main factor that determines the size of a species’ ecological niche is the availability of resources to individuals within the species.

Just as the competition can promote specialization, it can act antagonistically towards specialization and promote generalization. It is hypothesized that greater intraspecific competition (usually by way of a larger population) leads to a species’ generalization (referred to as the competitive diversification hypothesis by Jones and Post (2016)) (Araújo et al., 2011; Jones and Post, 2013; Jones and Post, 2016). According to the hypothesis, if the individuals within a population are alike or broadly similar in niche preference, then more individuals will lower fitness in core niche space (usually due to a decline in resource availability). As fitness in core niche space declines, individuals who capitalize on the marginal niche spaces gain a relative fitness advantage leading to a diversification of resource use and overall generalization within the population (Fig. 4). While intuitive, non-theoretical studies have revealed mixed results with some showing greater specialization with increased population size and more intraspecific competition (reviewed in Jones and Post, 2016). Jones and Post (2013; 2016) developed their own hypothesis that whether a population generalizes or specializes depends upon the strength of competition with higher levels of competition leading to specialization (Fig. 4). This hypothesis called the intermediate competitive diversification hypothesis extends the competitive diversification hypothesis and offers a more nuanced look at the interaction between niche width and intraspecific competition.

Previous theoretical studies have looked how individuals change their niche position, and its effect on the population’s niche, with increasing population (Roughgarden, 1972; Svanback and Bolnick, 2005; Abrams et al., 2008). That said, individuals do not use a single resource and can be flexible and use other resources; in this way, individuals themselves also have a niche width. Population niche width will change with individual niche width, and such changes can be nearly identical if individual variation in niche position is low (Bolnick et al., 2003). We seek to understand how individual niche width may change with increasing population size and what effect this has on the entire population. To that end, we created and analyzed a game theoretic model of ecological specialization inspired by Ackermann and Doebeli (2004). We used it to assess if an increased population and intraspecific competition lead to increased specialization. In our model, populations change niche position and specialization in response to various biotic and abiotic conditions. Using this model, we performed a parameter sweep to see how a population’s optimal niche space varied in response to population size under differing resource spreads and penalties to specialization. We show that increased population specialization can happen with increased population density especially when the population size is low, resources are compact, and the penalty to specialization is low. We hypothesize that this is due to a competitive advantage of specialization, adding to previous hypotheses on the interaction between niche width and intraspecific competition.

## Methods

In order to understand competitive diversification, we created a two-trait, evolutionary game theoretic model of resource use based on the G-function developed by Vincent and Brown (1987, 2005). In this notation, the population growth rate of species *i* is defined as

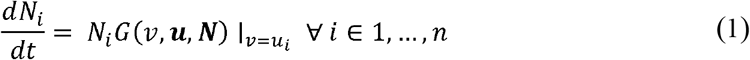

where *G* (*ν*, ***u***, ***N***) is the fitness (as defined by per-capita growth rate) of a focal individual with strategy *ν* in a community, where ***u*** = (*u*_1_, … , *u_s_*) is the vector of strategies found among the *s* species in the community and ***N*** = (*N*_1_, … , *N_s_*) is the vector of population densities for each of the *s* species. This fitness generating function *G* (*ν*, ***u***, ***N***) generates the fitness of species *i* when *ν* is set equal to *u_i_*. Taking the derivative of the fitness generating function with respect to *ν* and further substituting *ν* for *u_i_* gives us the evolutionary dynamics for species *i*

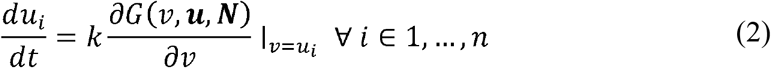

where *k* some is measure of additive genetic variance for natural selection.

Our model follows the concept of the resource utilization curve introduced by MacArthur (1972) and is inspired by Ackermann and Doebeli (2004). First, we assume that there exists some resource in the environment available to all species. This resource may vary along one or more continuous or discrete attribute axes. One can imagine the resource being seeds and an attribute being seed size. We approximate the distribution of resource abundances based on attributes as Gaussian (equation 3) (Fig. 1a).

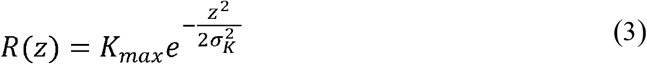

**Figure 1.**
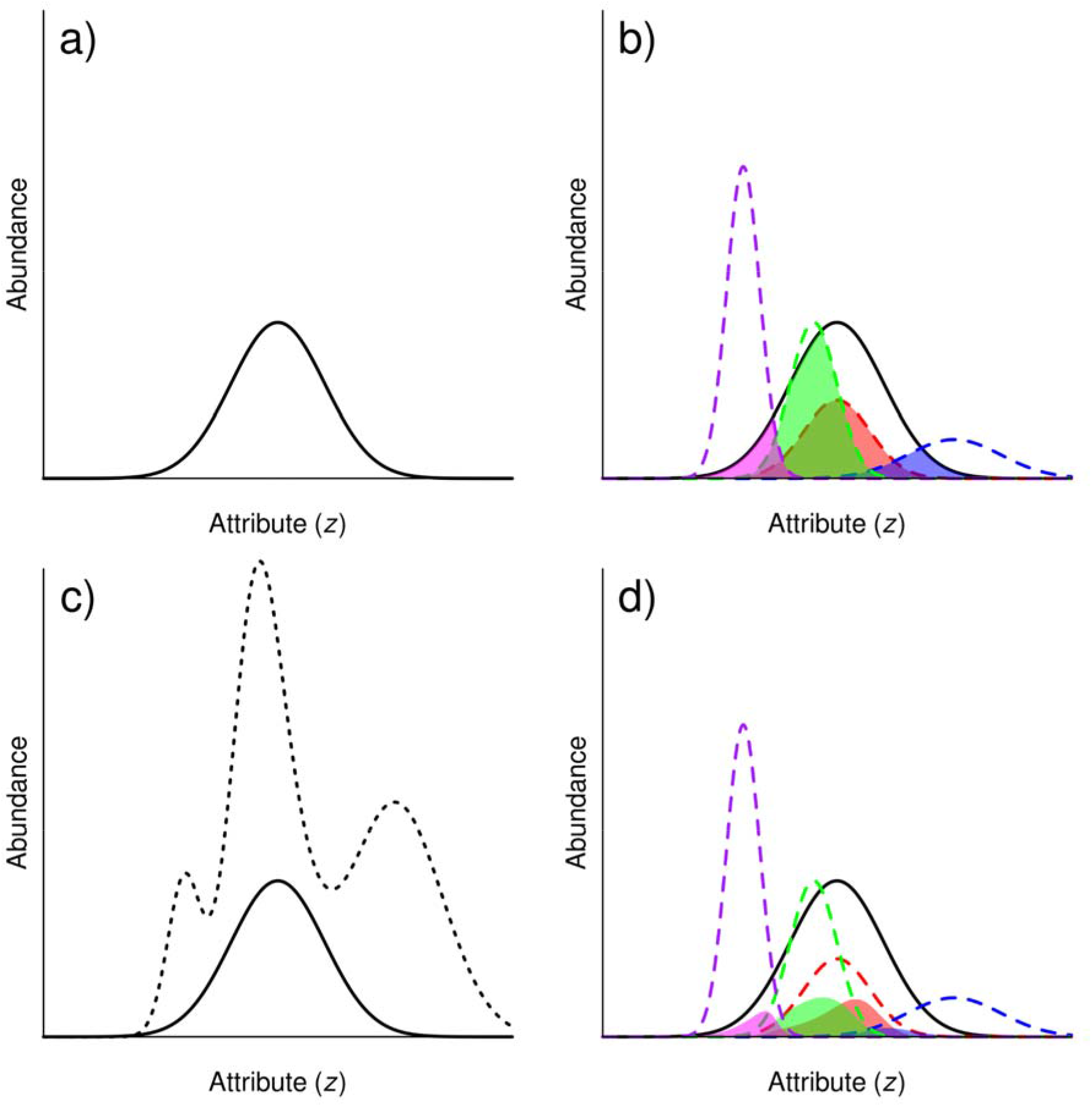
A visual representation of our mathematical model. (a) Abundance of a resource given its attribute. We assume the resources are so finely divided and packed together that the resource class is taken to the limit and represents a continuous Gaussian distribution (solid line). (b) The individual utilization curves of four different species, here represented by different colors. Each utilization curve determines how well an individual can gather and assimilate the resource assuming there are enough resource (dashed lines). Individual utilization curves vary in preference (niche position) and flexibility (width) which alters the total amount of resources the respective individual can use. The actual amount captured by each individual in the absence of competition depends upon the abundance of resources and is represented by the shaded areas. (c) The utilization curves of individuals from many different species can be added together to grant the total utilization curve for the community. (d) Individuals are forced to share the resource shrinking the actual amount of resource captured by each individual (the new shaded area).

Here, *R(z)* specifies the abundance of the resource (seed) of a particular attribute *z* (size). In our model, we assume that the attribute *z* is discrete. The resource with the highest abundance has the attribute *z* = 0, and abundances fall off according to the rate *σ*_K_. In this way, *σ*_K_ determines the spread of the resource attributes. A smaller *σ*_K_ denotes a more rapid decline in resource availability as the attributes of the resource deviates from *z* = 0. In our example,*z* = 0 may represent medium size seeds (on a log scale) while *σ*_K_ determines the number of small and large seeds relative to medium seeds. Resources are assumed to have a high replenishment rate leading to timescale separation between the resource and species dynamics and a fixed amount of resources in the environment.

With this resource distribution, we envision how an individual organism may capture and utilize resources of various attributes. Equation 4 describes how much of a resource an individual can capture at a given instant in time – its utilization curve. We assume that the utilization curve is also Gaussian and is maximized at a resource of a specific attribute (Fig. 1b).

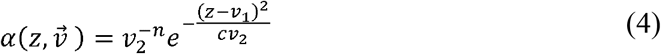

This utilization curve determines the amount of the resource of attribute *z* that can be captured by an individual based on its suite of microevolutionary adaptations 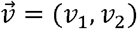. In this model, *ν*_1_ represents the type of resource the individual is most efficient at capturing (specifically when *z* = *ν*_1_) and can be thought of as the niche position. As well, *ν*_2_ determines how efficiently it captures resources different from *z* ≠ *ν*_1_ and can be thought of as the inverse of specialization (larger *ν*_2_ means a less specialized individual). We can think of *ν*_1_ as the preferred resource for the individual and *ν*_2_ as its flexibility in resource use. *ν*_1_ does not fundamentally alter the curve, it merely shifts it; *ν*_2_, on the other hand, does. As specialization increases *ν*_2_ → 0, the total amount of resources captured around *ν*_1_ – an organism’s core resources – increases while capture rate of those farther away from *ν*_1_ – the marginal resources – decreases. One can imagine the organism to be a seed-eating bird. For this bird, *ν*_2_ may denote how flexible and effective it is consuming seeds of various sizes while *ν*_1_ may denote the behavioral preference of the bird for a specific seed size.

Under the G-function framework, 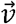 represents the microevolutionary adaptations of an organism. These traits evolve on the order of population timescales. The other parameters (*c* and *n*) represent macroevolutionary adaptations which evolves on timescales orders of magnitude larger than population timescales and can be considered as relatively fixed. These are the fundamental constraints that govern the individual’s utilization of the resource. Parameter *c* can be thought of as the intrinsic constraints in how efficiently an organism can gather multiple different resources with a larger *c* denoting fewer constraints. Here, *c* is a strictly positive parameter that determines the tradeoff between obtaining marginal resources versus core resources with changes in specialization *ν*_2_. If 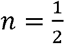, the individual will gain as much in resources as it loses with increasing specialization; if 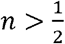, then the gain with increasing specialization is greater than the loss; and if 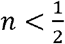, the loss is greater than the gain. We call *n* the penalty to specialization.

Substituting 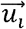 for 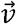, we get the utilization curve of an individual of species *i*. Assuming individuals of the same species are identical in 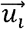, we can get a utilization curve of the entire species 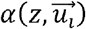 where *N_i_* is the population density of the species. If we sum of all utilization curves (including the utilization of the focal individual), we can get a “total utilization curve” for the community (equation 5) (Fig. 1c).

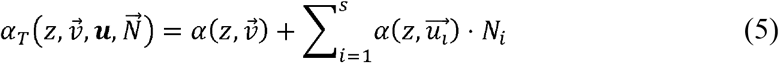

Here, 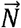 is a vector of population density among *s* different species, **u** is a 2 × *s* matrix with the strategies of the species, and 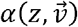 remains the utilization curve of a focal individual. If the amount of resources at attribute *z* are sufficient for the entire community 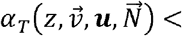 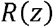 then an individual’s capture rate at that attribute is simply 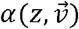. If however, the amount of resources is insufficient for the entire community 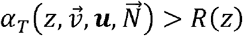, then competition occurs and we assume the individual must “share” those resources with the community. In such a case, the amount of resources the individual captures is proportionate to the total amount of resources. This gives us the actual utilization curve (equation 6) (Fig. 1d).

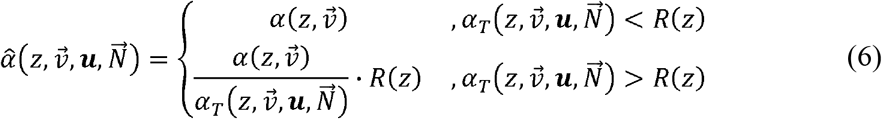

Summing the amount of each resource captured by an individual over all resources gives the total amount of resources by an individual (equation 7).

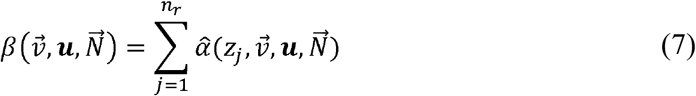

We assume all resources captured by an individual is converted into reproduction and the creation of more individuals. We also assume that the species’ death rate *d* is density independent, giving us the full G-function.

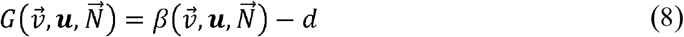

In order to test the hypothesis of competitive diversification, we saw how optimal specialization 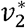 changed with population density *N* under different parameter sets. We assumed a fine but discrete class of resources. In our seed example, each seed can be lumped into categories of discrete size but transition between each seed size is so fine that it is essentially imperceptible to the organisms. This division of the resource class is taken to the limit such that it becomes a continuum (see SI). We assumed there was only intraspecific competition, no interspecific competition, giving us a single species with identical traits. If the resources are symmetrical about 0, then *ν*_1_ is always optimized at 0 for the population regardless of other parameters. Therefore, only *ν*_2_ need be optimized (see SI). To determine optimal specialization, we fixed all parameters including species density and solved for the *ν*_2_ such that

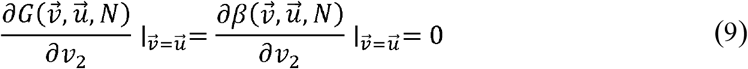

Since optimal specialization cannot be solved analytically, we analyzed it numerically. Using this basic setup, we varied the population density *N* from 0 to 10 in increments of 0.01 and saw how 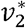 changed in response to population size. We did this under varying resource spread *σ_K_* ranging from 0.1 to 10 by increments of 0.1 and penalty to specialization *n* ranging from 0.05 to 5 by increments of 0.05 to see the influence of resource spread and penalty to specialization affects competitive diversification. We fixed parameters *K_max_* = 1 and *c* = 2.

## Results

With the parameters selected, we see a wide variation in optimal specialization values, ranging from highly specialized 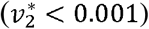 to highly generalized at 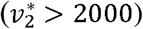. Presenting this on a log scale, the range is largely symmetrical ranging from *ν*_2_ = −6.32 to *ν*_2_ = 7.69. The median value of optimal specialization is *ν*_2_ = −0.62with the majority being centered at that value (50.81% lie within the range −1 to 0 and 70.72% lie within the range −1.5 to 0.5). With this basic analysis, we feel confident to have selected a broad enough range in parameters to cover a broad range of model behavior.

Looking at the results generally, we can say that optimal specialization increases (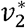 decreases) with lower resource spread and a lower penalty to specialization (Fig. 2c, 3a). This makes sense as a lower resource spread reduces available choices, making specialization on the most abundant more advantageous. In fact, the effects of spread appear non-linear with the most extreme specialization values occurring with *σ_K_* ≤ 1 for a given penalty to specialization and population density (Fig. 2a,b,c). As well, a lower penalty to specialization means less of a tradeoff which makes specialization more advantageous.

**Figure 2.**
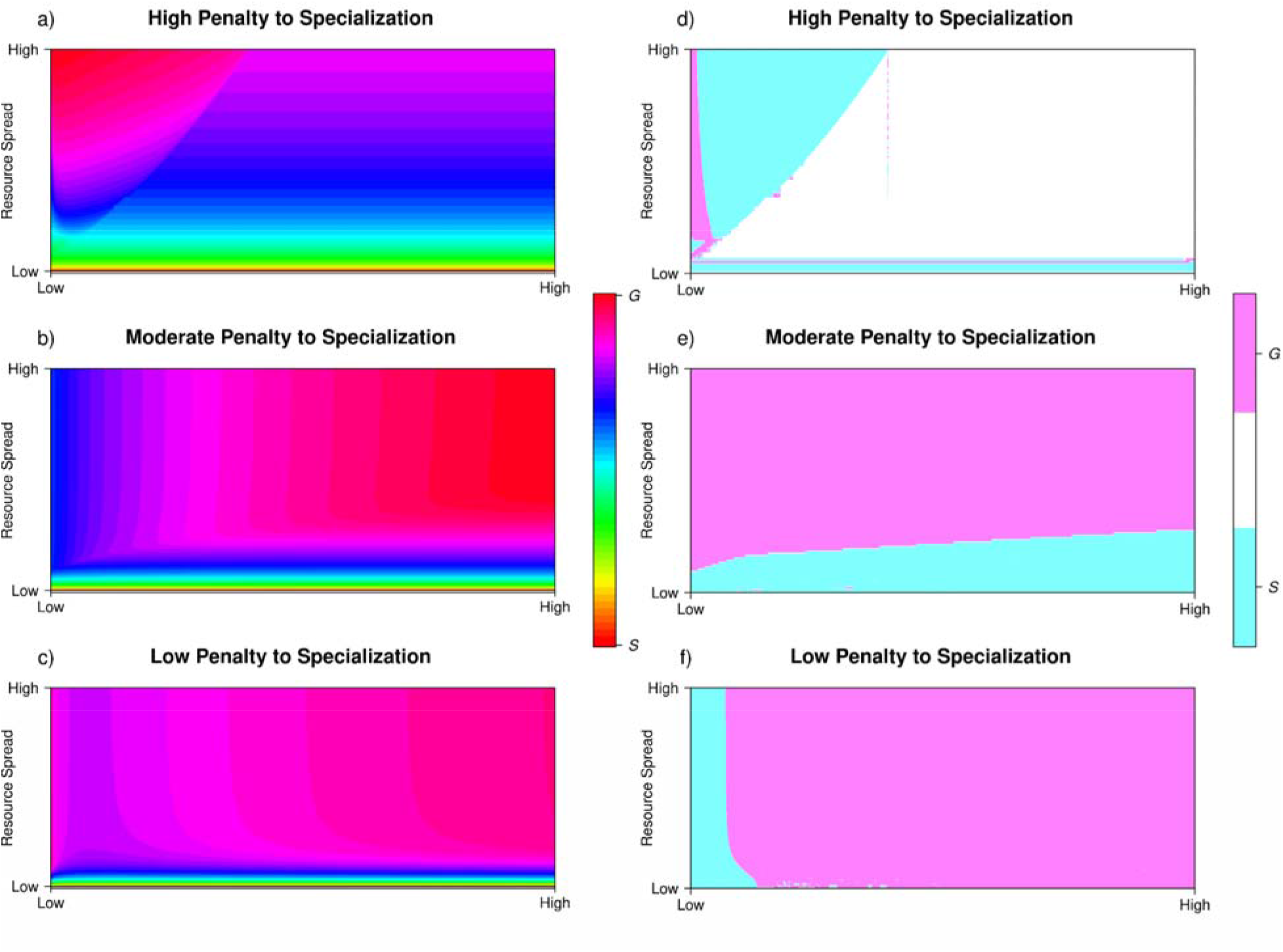
(a, b, c) Optimal niche width with varying population densities and resource spread given a penalty to specialization. Warm colors indicated a population with a smaller niche width (more specialized) while cool colors indicated a population with a larger niche width (more generalized). The data presented are log transformed. (d, e, f) How niche width changes with resource spread and incremental increases in population. This figure shows the sign of the difference in niche width between adjacent population sizes. Cyan colors indicate a shrinking niche width (specialization) with an incremental increase in the population while magenta colors indicate the opposite. Our results show the presence of increasing specialization with increasing population size under a variety of conditions. This can occur like the intermediate competitive diversification hypothesis with a decrease in specialization before an increase in specialization (d, e), as a persistent increase in specialization which seems to be more common with a low spread of resources (d, e), or at low population sizes before increased generalization as seen with a low penalty to specialization (f).

**Figure 3.**
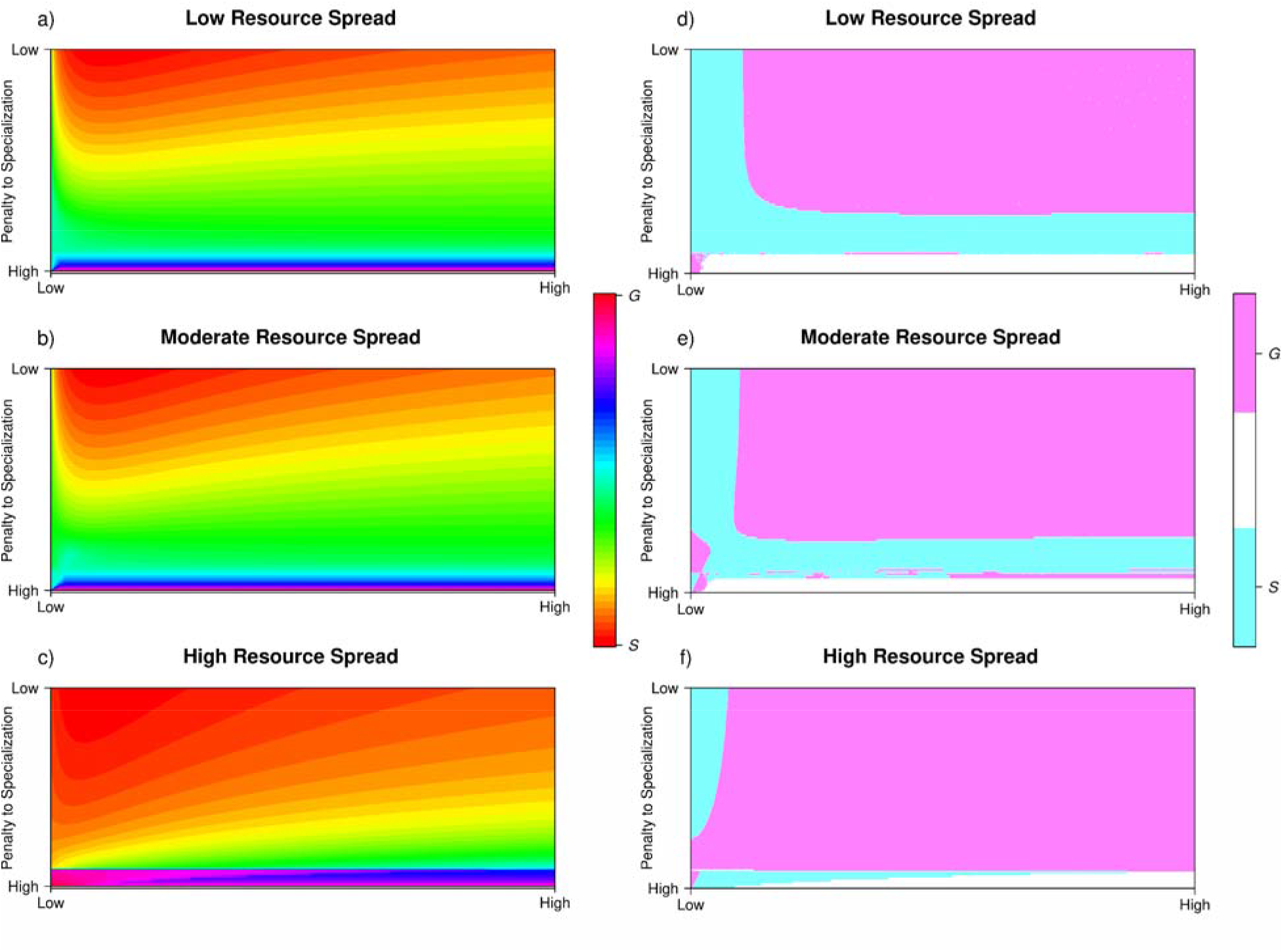
(a, b, c) Optimal niche width with varying population densities and penalty to specialization given a resource spread. (d, e, f) How niche width changes with resource spread and incremental increases in population. The colors indicate the same phenomena as in Figure 2. Our results show the presence of increasing specialization with increasing population size under a variety of conditions. This can occur like the intermediate competitive diversification hypothesis with a decrease in specialization before an increase in specialization (d, e), as a persistent increase in specialization which seems to be more common with a low spread of resources (d, e), or at low population sizes before increased generalization as seen with a low penalty to specialization (f).

With regard to population, increased population density can increase optimal specialization, but the specifics depend on the parameters. With a moderate penalty to specialization and high resource spread, we see what is assumed under the competitive diversification hypothesis – increasing the population size increases the niche width of the population (Fig. 2e, 3f). If resource spread is moderate and the penalty to specialization is moderate, then we see what we expect from the intermediate competitive diversification hypothesis, namely an increase in the niche width followed by a decrease (Fig. 2e,3e). We also see this phenomenon when resource spread is fairly high and the penalty to specialization is high (Fig. 2d,3f).

We also see some new phenomena not captured by either hypotheses. Firstly, there are areas which show a constant increase in specialization. This occurs with low resource spread and a moderate penalty to specialization as well as a high penalty to specialization (Fig. 2d,3d). We also sometimes see an increase in specialization follow by generalization. This mostly occurs with a low penalty to specialization, regardless of how much spread there is in the resources, and a little bit when there is a high penalty to specialization and resource spread is moderate (Fig. 2d,2f,3e).

## Discussion

Intraspecific competition is generally thought to lead to a generalization in resource use, but recent evidence suggests that the opposite can happen as equally as likely (Jones and Post, 2016). We sought to examine the conditions under which one might see this response by creating and analyzing a game-theoretic model of resource use. Our analysis shows that increasing population density (and thereby increasing intraspecific competition) can lead to greater specialization of the population. This especially occurs at lower absolute population densities. Other conditions likely to promote increased specialization are both a low penalty to specialization and resource spread. Our model adds to the literature by showing that increased specialization with increased competition can happen at low levels of competition.

Some of our results concur with previous work. When the penalty to specialization and resource spread are high, specialization decreases before increasing with respect to population (Fig. 2d,3f). This pattern replicates the expectation of the intermediate competitive diversification hypothesis which states that moderate levels of competition lead to a generalization but high levels of competition lead to specialization as resources become ever more depleted (Fig. 4) (Jones and Post, 2016). We also see high specialization and convergence when resource spread is low (Fig. 3a). Previous studies have shown that strategy convergence can occur when resources are essential (Abrams, 1987; Fox and Vasseur, 2008). While our resources are substitutable, the lack of available options means that when resource spread is low, the central resource is essentially essential.

**Figure 4.**
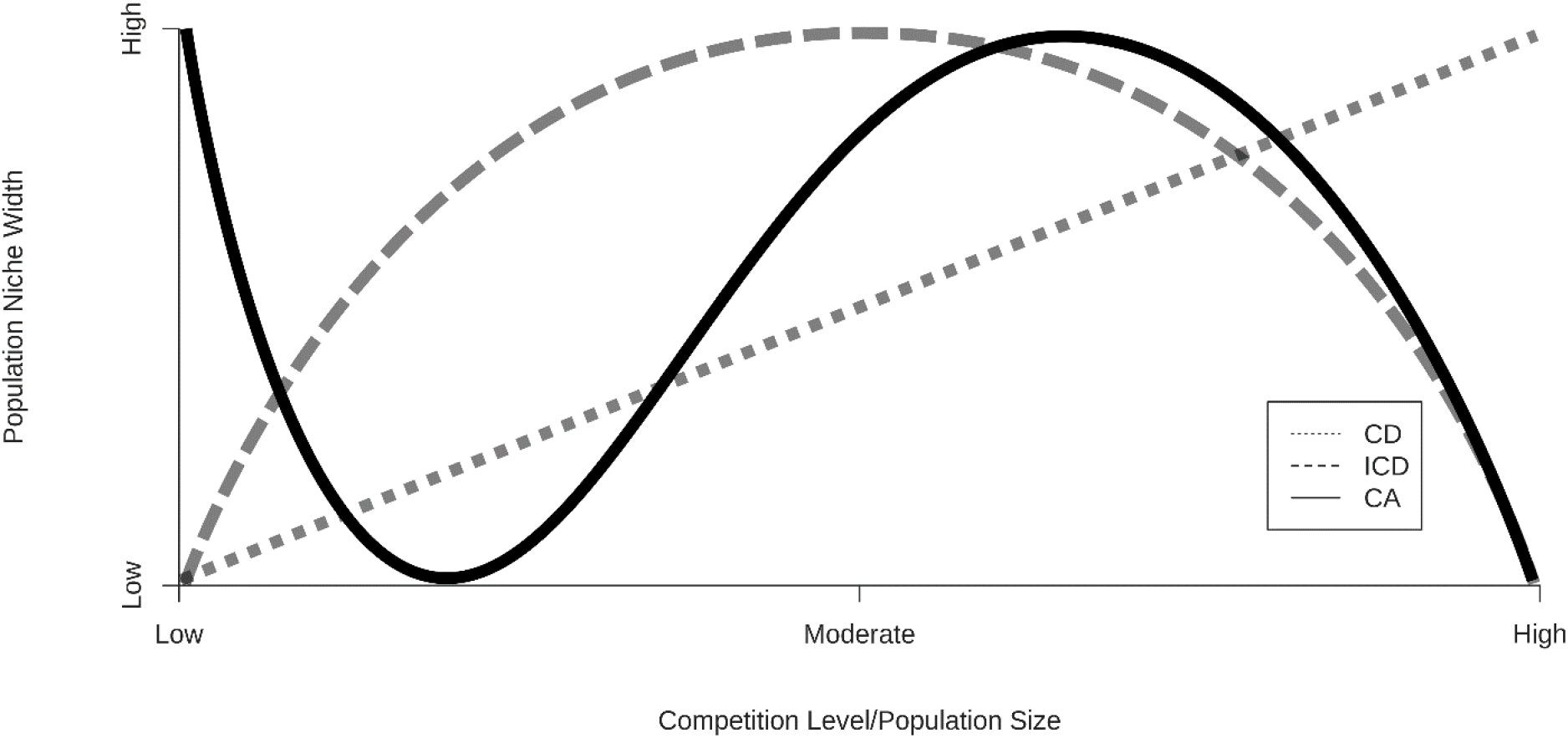
A schematic comparing our results to previous hypotheses. Under the competitive diversification hypothesis (CD), increases in the level of competition and population size lead to increased population niche width regardless (dotted line). Under the intermediate competitive diversification hypothesis (ICD), increased intraspecific competition at low levels results in greater population niche width but increases at high levels result in a smaller population niche width (dashed line). Our results show that indicate that there may be a competitive advantage to specialization such that increasing competition can increase specialization which seems to primarily occur at lower population sizes (CA, solid line).

Our model also suggests new results. One thing we see is multiple points of optimal specialization (Fig. S1). This particularly occurs when resource spread is high and the penalty to specialization is high or moderate. We are currently unsure why this occurs, why it is restricted to these areas, or what it may mean. As mentioned earlier, we also see that increased specialization can be the initial response to an increased population, especially when the penalty to specialization is low. This creates a U-shaped response of population niche width to population density, opposite of the hump-shaped response of the intermediate competitive diversification hypothesis (Fig. 4). We reckon that this may be due to a competitive advantage that comes with specialization. Previous studies have shown that specialist species can be more competitive than generalists in the competition for resources, but have looked at it primarily from an interspecific framework (MacArthur and Levins, 1964; Dykhuizen and Davies, 1980). Our study seems to suggest that specialization can offer a competitive advantage within species and populations. Under our model, if there is an insufficient amount of a resource, then that resource will be shared amongst species within the community proportional to each species utilization of said resource. Those more specialized on a resource will have higher utilization rates which means they get proportionally more of the resource and therefore could potentially have higher fitness. This creates a competitive advantage to specialization, especially when the penalty is low. We can also see this competitive advantage when looking at the non-linear response of specialization to resource spread. Even though resource spread declines linearly, specialization increases rapidly (Fig. 2a,b,c). This non-linear response may occur because competition becomes increasingly intense as resources become scarcer.

Our results may show this because we analyzed for the niche width of individuals and not just niche position. Previous analyses looked at how a population changes its niche width by looking at how individuals change their niche position and preferred resources with competition (Roughgarden, 1972; Abrams et al., 2008; Ackermann and Doebeli, 2004). Our analysis instead directly measured the change of individual niche width with respect to population size while individuals of the population kept the same resource preference (Rosenzweig, 1991). Having the same resource preference enhances the effects of competition and therefore increases the advantage specialization brings. This disconnect is also borne out in our analysis. We also determined whether the population given all the parameters showed stabilizing or disruptive selection on niche width by calculating whether the population is at a maximum or minimum of the adaptive landscape respectively with respect to niche position 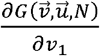. Firstly, disruptive vs. stabilizing selection seems to correlate with absolute specialization level. Secondly, with increasing population, more often there is solely disruptive selection, solely stabilizing selection, or disruptive followed stabilizing selection with increasing population size (Fig. S2, S3). Only in a few cases, namely with high resource spread and low to moderate penalty to specialization, do we see stabilizing followed by disruptive selection, and even so it is only at higher population. At lower population sizes under those conditions, there is disruptive followed by stabilizing selection (Fig. 4, S2b,c, S3c). These differences are likely the reasons as to why our study revealed the competitive advantage of specialization.

Through our study, we have generated a hypothesis on the reason as to why increased population specialization is seen with increased competition in nature. Whether or not it is valid to a particular natural system depends on several things. Firstly, when competition increases and preferred resources deplete, an individual has two options: shift resource preference or broaden resource use with the former leading to more individual variation (Rosenzweig, 1991; Svanback and Bolnick, 2005). If the majority of organisms retain a shared preference for a resource (as it is more abundant or calorific) and intraspecific variation is minimal, then individuals broadening their resource use may be the more immediate response to changes in population specialization. Individual variation in resource in populations use can be high or low and is overall largely equivocal. Secondly, shifts in preferred resource versus a broadening of resource use may be a function of whether they are a behavioral vs morphological response. For example, Smith (1990) showed that though large-billed and small-billed varieties of an African finch, *Pyrenestes ostrinus*, shared similar preferences for soft seeds, the large-billed individuals were able to more readily switch to harder seeds when the softer seeds were rarer. In this case, shifts in resource preference were largely down to a behavioral response while the ability to broaden resource use was down to a morphological response (bill size). Determining the type of response based on the shift will be key to determining the changes in population specialization. Lastly, the initial population size/density from its increase remains an important factor. With competitive advantage, increased specialization with increased competition occurred when population (and therefore competition) was initially low while increased specialization with increased population occurred when competition was initially moderate to high for the intermediate competitive diversification hypothesis (Fig. 4) (Abrams et al., 2008; Jones and Post, 2013; Jones and Post, 2016). Determining the population size and the strength of competition will govern whether increased specialization was due to competitive advantage or competitive diversification. In our model, mortality is a strategy-independent term that governs the equilibrial population size. In this case, the competitive advantage reason may be seen among species with higher mortalities while the intermediate competitive diversification may be seen among species with lower mortalities. Taking into account these three factors would help tease apart the reasons for increased specialization.

We have created and analyzed a model of resource use with evolvable individual specialization to see how changes in population size affect the optimal specialization. We show that increases in the population, though generally lead to more generalized resource use, can lead to increases in specialization, particularly at low population sizes. We hypothesize that this may be due to the competitive advantage that specialization can bring. This work adds dimensions and flavor to previous work and offers a potential hypothesis as to why increased specialization may be seen in nature.

## Notes

### Competing Interest Statement

The authors have declared no competing interest.

