## Supplementary Information for "Intraspecific Competition and the Promotion of Ecological Specialization"

**Figure S1** A plot of $\frac{\partial G}{\partial v_{2}}$ vs.$v_{2}$ for various population densities $\left( N \right)$. As $N$ increases, we move from having only one optimal specialization point $\left( N=1 \right)$ to having three points of optimal specialization $\left( N=1.625, 2.25, 2.875, 3.5, 4.125 \right)$ to a maximum of five $\left( N=4.75 \right)$ back down to three $\left( N=5.375 \right)$ then one $\left( N=6 \right)$. Discontinuities in optimal specialization vs. population size invariably occur; the largest root for the population range of $N=1$ to $N=4.75$ seems to indicate continuity of optimal specialization but goes away at $N=5.375$ and $N=6$ leading to a jump in optimal specialization.

**Figure S2** A plot of selection on niche position $\frac{\partial^{2}G}{\partial v_{1}^{2}}$ for various levels of penalty to specialization $n$. Grey indicates positive values, and therefore disruptive selection on niche position and a divergence in resource preference, while green indicates negative values and therefore stabilizing selection on resource preference. Brown indicated a value of 0 and therefore neither disruptive nor stabilizing selection. Convergence and divergence in resource preference seem to correlate with the niche width of the population where a low niche width correlates with convergence in resource preference and a high niche width correlates with divergence in resource preference.

**Figure S3** A plot of selection on niche position $\frac{\partial^{2}G}{\partial v_{1}^{2}}$ for various levels of resource spread $\sigma_{K}$. The colors represent the same thing as Figure S2, and correlation between convergence/divergence and niche width remains the same.

Fig. 1

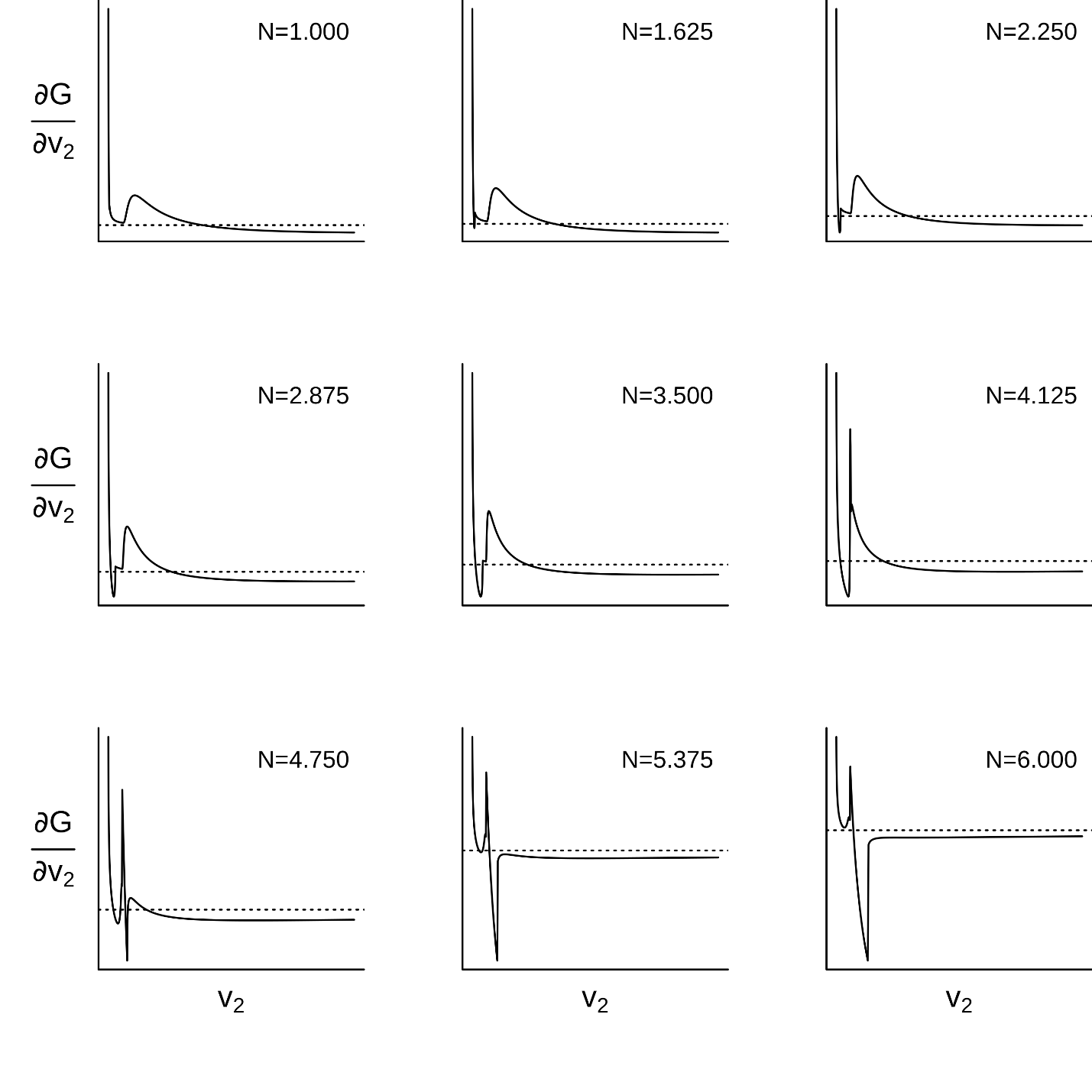

Fig. 2

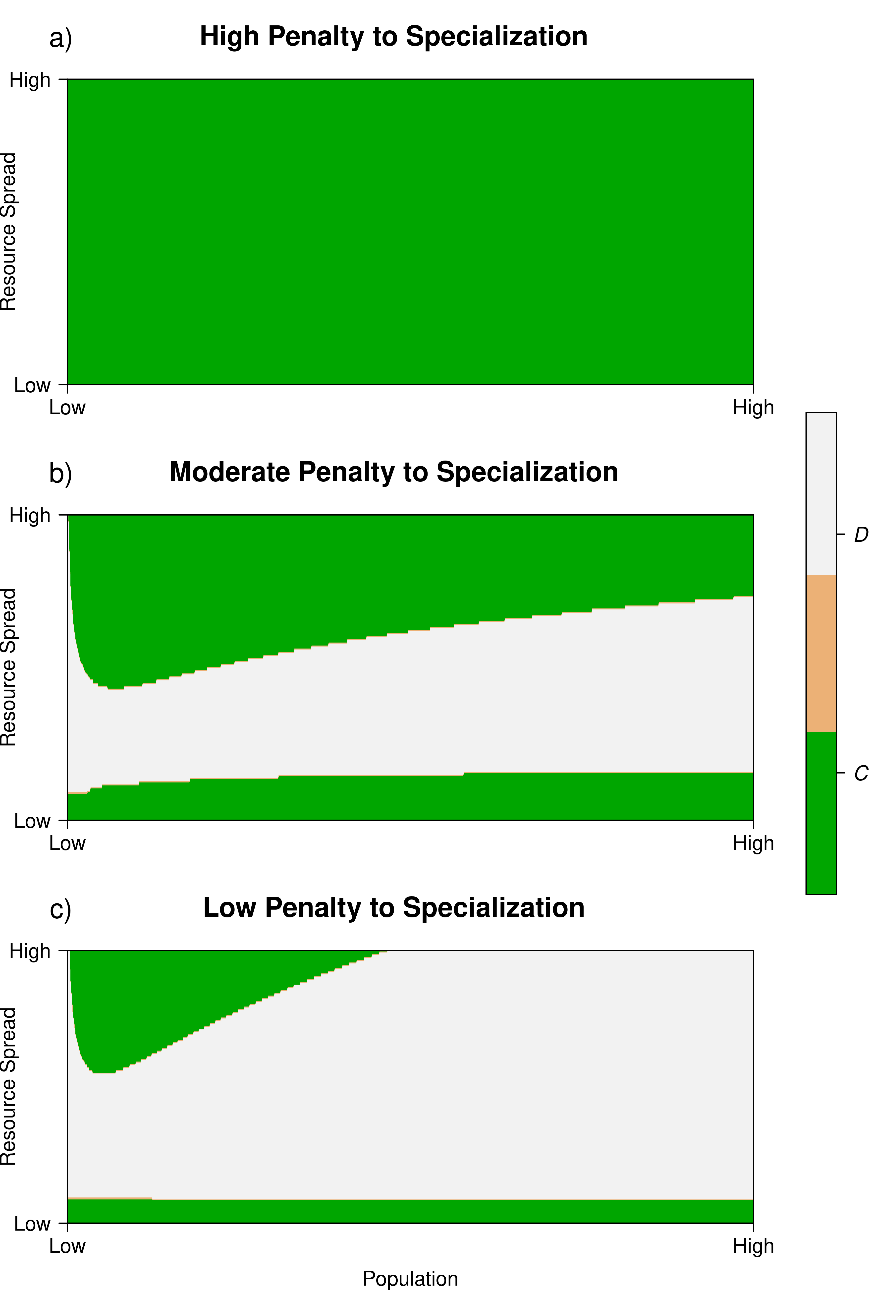

Fig. 3

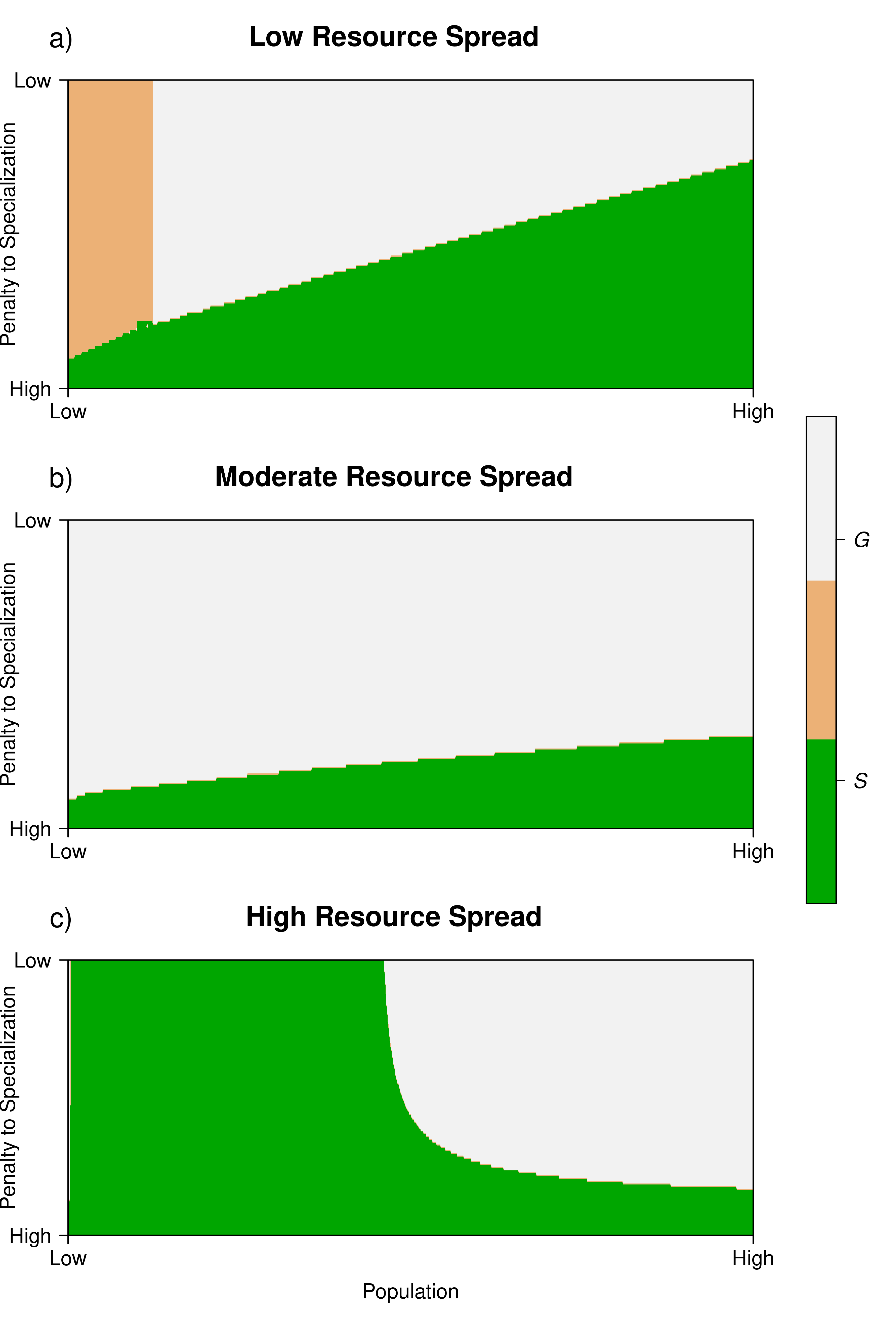

### **The G-Function Framework**

We present a brief overview of the G-function framework used for this model. In this framework, the population growth rate of species $i$ is defined as

$$\frac{dN_{i}}{dt}= N_{i}G\left( v,\boldsymbol{u},\boldsymbol{N} \right)\mid_{v=u_{i}} \forall i\in1,\ldots,n$$

(1)

where $G\left( v,\boldsymbol{u,N} \right)$ is the fitness (as defined by per-capita growth rate) of a focal individual with strategy $v$ in a community, where $\boldsymbol{u} = (u_{1},\ldots,u_{s})$ is the vector of strategies found among the $s$ species in the community and $\boldsymbol{N}= (N_{1},\ldots,N_{s})$ is the vector of population densities for each of the $s$ species. This fitness generating function $G\left( v,\boldsymbol{u},\boldsymbol{N} \right)$ generates the fitness of species $i$ when $v$ is set equal to $u_{i}$. Taking the derivative of the fitness generating function with respect to $v$ and further substituting $v$ for $u_{i}$ gives us the evolutionary dynamics for species $i$

$$\frac{du_{i}}{dt}=k\frac{\partial G\left( v,\boldsymbol{u},\boldsymbol{N} \right)}{\partial v}\mid_{v=u_{i}} \forall i\in1,\ldots,n$$

(2)

where $k$ is some measure of additive genetic variance for natural selection.

### **Calculating Optimal Specialization**

Our G-function is represented by the formula $G(\vec{v},\mathbf{u},\vec{N})\mid_{\vec{v}=\vec{u_{i}}}=\beta(\vec{v},\mathbf{u},\vec{N})\mid_{\vec{v}=\vec{u_{i}}}-d$ where $\beta(\vec{v},\mathbf{u},\vec{N})\mid_{\vec{v}=\vec{u_{i}}}=\sum_{j=1}^{n_{R}} \hat{\alpha}(z_{j},\vec{v},\mathbf{u},\vec{N})\mid_{\vec{v}=\vec{u_{i}}}$ and where

$$\hat{\alpha}(z_{j},\vec{v},\mathbf{u},\vec{N})=\left\{ \begin{matrix} \alpha(z_{j},\vec{v}) & ,\alpha_{T}(z_{j},\vec{v},\mathbf{u},\vec{N})<R(z_{j}) \\ \frac{\alpha(z_{j},\vec{v})}{\alpha_{T}(z_{j},\vec{v},\mathbf{u},\vec{N})}\cdot R(z_{j}) & ,\alpha_{T}(z_{j},\vec{v},\mathbf{u},\vec{N})>R(z_{j}) \end{matrix} \right.$$

(1)

In our case, $R\left( z \right)=K_{max}e^{-\frac{z_{j}^{2}}{2\sigma_{K}^{2}}}$, $\alpha\left( z_{j},\vec{v} \right)=v_{2}^{-n}e^{\frac{-(z_{j}-v_{1})^{2}}{cv_{2}}}$, and $\alpha_{T}(z_{j},\vec{v},\mathbf{u},\vec{N})=\alpha(z_{j},\vec{v})+\sum_{i=1}^{n_{s}} \alpha(z_{j},\vec{u_{i}})N_{i}$. Since each resource is independent of the other, we can say that $\frac{du_{2}}{dt}=\frac{\partial G(\vec{v},\mathbf{u},\vec{N})}{\partial v_{2}}\mid_{\vec{v}=\vec{u_{i}}}=\frac{\partial\beta(\vec{v},\mathbf{u},\vec{N})}{\partial v_{2}}\mid_{\vec{v}=\vec{u_{i}}}=\sum_{j=1}^{n_{R}} \frac{\partial\hat{\alpha}(z_{j},\vec{v},\mathbf{u},\vec{N})}{\partial v_{2}}\mid_{\vec{v}=\vec{u_{i}}}$ where

$$\frac{\partial\hat{\alpha}(z_{j},\vec{v},\mathbf{u},\vec{N})}{\partial v_{2}}=\left\{ \begin{matrix} \frac{\partial\alpha(z_{j},\vec{v})}{\partial v_{2}} & ,\alpha_{T}(z_{j},\vec{v},\mathbf{u},\vec{N})<R(z_{j}) \\ \frac{\partial\alpha(z_{j},\vec{v})}{\partial v_{2}}\cdot\frac{\sum_{i=1}^{n_{s}} \alpha(z_{j},\vec{u_{i}})N_{i}}{\alpha_{T}(z_{j},\vec{v},\mathbf{u},\vec{N})^{2}}\cdot R(z_{j}) & ,\alpha_{T}(z_{j},\vec{v},\mathbf{u},\vec{N})>R(z_{j}) \end{matrix} \right.$$

(2)

with $\frac{\partial\alpha(z_{j},\vec{v})}{\partial v_{2}}=\frac{((z-v_{1})^{2}-cnv_{2})}{cv_{2}^{2}}\alpha(z_{j},\vec{v})$

As the number of resources $n_{R}$ increases, $\beta(\vec{v},\mathbf{u},\vec{N})\mid_{\vec{v}=\vec{u_{i}}}$will go to infinity, causing similar extremes for $\frac{\partial\beta(\vec{u},\mathbf{u},\vec{N})}{\partial v_{2}}\mid_{\vec{v}=\vec{u_{i}}}$. If the number of resources is not increasing but instead being more finely divided, we can multiply each interaction by a decreasing weight$w$such that$\lim_{n_{R}\to\infty}w=0$. We arrive at a new equation$\beta(\vec{v},\mathbf{u},\vec{N})\mid_{\vec{v}=\vec{u_{i}}}=\sum_{j=1}^{n_{R}} w_{j}\cdot\hat{\alpha}(z_{j},\vec{v},\mathbf{u},\vec{N})\mid_{\vec{v}=\vec{u_{i}}}$. Similarly, $\frac{\partial\beta(\vec{u},\mathbf{u},\vec{N})}{\partial v_{2}}\mid_{\vec{v}=\vec{u_{i}}}=\sum_{j=1}^{n_{R}} w_{j}\cdot\frac{\partial\hat{\alpha}(z_{j},\vec{v},\mathbf{u},\vec{N})}{\partial v_{2}}\mid_{\vec{v}=\vec{u_{i}}}$. Assuming $w_{j}$is constant ($w_{j}=w$ for all$j=1,2,...,n_{R}$), it simply gets folded into the speed of the evolutionary and population dynamics and does notaffect the equilibrium. This weight instead acts like a Reimann summation and therefore, the species’ interactions with an infinitely fine discrete resource base can be calculated as an integral $\frac{du_{2}}{dt}=\int_{-\infty}^{\infty} \frac{\partial\hat{\alpha}(z,\vec{v},\mathbf{u},\vec{N})}{\partial v_{2}}\mid_{\vec{v}=\vec{u_{i}}}dz$.

To obtain full evolutionary dynamics, we must obtain the values of the resource attribute $z=\hat{z}$such that $\alpha_{T}(\hat{z},\vec{v},\mathbf{u},\vec{N})=R(\hat{z})$ as the function is piecewise. Because $\alpha_{T}$ is the sum of multiple exponential functions and $R(z)$ is another exponential function, we cannot arrive at a general solution for evolutionary dynamics. Instead, we can arrive at a more limited solution using some key assumptions. Firstly, we can assume that there is only one population using a single strategy $\vec{u}$(or all species use the same srategy). Secondly, we only look to derive evolutionary dynamics whe$n \vec{v}=\vec{u}.$Doing so means that we numerically solve for the equilibrium. Because of these two assumptions,$\alpha_{T}(z,\vec{v},\mathbf{u},\vec{N})\mid_{\vec{v}=\vec{u}}=(N+1)\alpha(z,\vec{u}).$ Therefore, we now have an exponential function equalling an exponential function, making the evolutionary dyanmics solveable. Since $R(z)$is symmetrical about 0, we can also assume that $u_{1}=0.$Altogether, solving for$\hat{z}$ yields

$$\hat{z}=\sqrt{\frac{-2\sigma_{K}^{2}cu_{2}log\left( \frac{K_{max}u_{2}^{n}}{N+1} \right)}{2\sigma_{K}^{2}-cu_{2}}}$$

(3)

Within the $\hat{z}$, there is the potential for the square root of a negative number. Because $\sigma_{K}^{2}cu_{2}$ is strictly positive, only $log\left( \frac{K_{max}u_{2}^{n}}{N+1} \right)$ and $2\sigma_{K}^{2}-cu_{2}$ can affect the sign of $\hat{z}$. Hereafter, we refer to these terms as $H$ and $S$ respectively. Looking at the terms, we can view them as comparing the resource function $R(z)$ and the total utilization function $\left( N+1 \right)\cdot\alpha(z,\vec{u})$. The first term $H=log\left( \frac{K_{max}\cdot u_{2}^{n}}{N+1} \right)$ compares the "height" of each curve at resource $z=0$. If positive, then the amount of resource $z=0$ is sufficient for all individuals $R\left( 0 \right)>\left( N+1 \right)\cdot\alpha(0,(0,u_{2}))$; if negative, then the amount is insufficient $R\left( 0 \right)<\left( N+1 \right)\cdot\alpha(0,(0,u_{2}))$. The second term compares $S=2\sigma_{K}^{2}-cu_{2}$ the "spread" of each curve. If positive, then at the furthest resources $\lim_{z\to-\infty,\infty}$ there will be sufficient resources for all individuals $\lim_{z\to-\infty,\infty}R\left( z \right)>\left( N+1 \right)\cdot\alpha(z,(0,u_{2}))$; if negative, then insufficient $\lim_{z\to-\infty,\infty}R\left( z \right)<\left( N+1 \right)\cdot\alpha(z,(0,u_{2}))$. For there to be a positive $\hat{z}$, the first term and the second term must be opposite signs. We can go on to break this down as such:

|  | $S>0$ | $S=0$ | $S<0$ |
| --- | --- | --- | --- |
| $H>0$ | Sufficient Amounts of All Resources | Sufficient Amounts of All Resources | Sufficient Amounts of Core Resources  Insufficient Amounts of Marginal Resources |
| $H=0$ | Sufficient Amounts of All Resources | Treated as Insufficient Amounts of All Resources | Insufficient Amounts of All Resources |
| $H<0$ | Insufficient Amounts of Core Resources  Sufficient Amounts of Marginal Resources | Insufficient Amounts of All Resources | Insufficient Amounts of All Resources |

Table S1: Comparison between resource curve and total utilization curve

Let $f_{1}=\frac{\partial\alpha(z,\vec{v},\mathbf{u},\vec{N})}{\partial v_{2}}\mid_{\vec{v}=\vec{u}}\mathrm{and}f_{2}=\left( \frac{\partial\alpha\left( z,\vec{v} \right)}{\partial v_{2}}\cdot\frac{\alpha\left( z,\vec{u} \right)\cdot N}{\alpha_{T}(z,\vec{v},\vec{u},N)^{2}} \right)\mid_{\vec{v}=\vec{u}}\cdot R(z)$. If there are sufficient amounts of all resources, then $\frac{du_{2}}{dt}=\int_{-\infty}^{\infty} f_{1}dz$. If there are insufficient amounts of all resources, then$\frac{du_{2}}{dt}=\int_{-\infty}^{\infty} f_{2}dz$. If there are sufficient amounts of core resources and insufficient amounts of marginal resources, then $\frac{du_{2}}{dt}=\int_{-\infty}^{-\hat{z}} f_{2}dz+\int_{-\hat{z}}^{\hat{z}} f_{1}dz+\int_{\hat{z}}^{\infty} f_{2}dz$. If there are insufficient amounts of core resources and sufficient amounts of marginal resources, then $\frac{du_{2}}{dt}=\int_{-\infty}^{-\hat{z}} f_{1}dz+\int_{-\hat{z}}^{\hat{z}} f_{2}dz+\int_{\hat{z}}^{\infty} f_{1}dz$. Solving for each component reveals

- $\int_{-\infty}^{\infty} f_{1}dz=\left( \frac{1}{2}-n \right)u_{2}^{-(n+1)}\sqrt{\pi cu_{2}}$
- $\int_{-\infty}^{\infty} f_{2}dz=\sigma_{K}\sqrt{2\pi}\frac{K_{max}\cdot N}{cu_{2}(N+1)^{2}}(\sigma_{K}^{2}-ncu_{2})$ ^𝑔𝑚𝑎_
- $\int_{-\hat{z}}^{\hat{z}} f_{1}dz=\left( \frac{1}{2}-n \right)u_{2}^{-(n+1)}\sqrt{\pi cu_{2}}\cdot erf\left( \frac{\hat{z}}{\sqrt{cu_{2}}} \right)-\hat{z}e^{\frac{-\hat{z}^{2}}{cu_{2}}}$
- $\int_{-\infty}^{-\hat{z}} f_{1}dz=\int_{\hat{z}}^{\infty} f_{1}dz=\frac{1}{2}\left( \left( \frac{1}{2}-n \right)u_{2}^{-\left( n+1 \right)}\sqrt{\pi cu_{2}}\cdot erfc\left( \frac{\hat{z}}{\sqrt{cu_{2}}} \right)-\hat{z}e^{\frac{-\hat{z}^{2}}{cu_{2}}} \right)$
- $\int_{-\hat{z}}^{\hat{z}} f_{2}dz =\sigma_{K}\sqrt{2\pi}\frac{K_{max}\cdot N}{cu_{2}(N+1)^{2}}(\sigma_{K}^{2}-ncu_{2})\cdot erf\left( \frac{\hat{z}}{\sqrt{2\sigma_{K}^{2}}} \right)-2\hat{z}\sigma_{K}^{2}e^{\frac{-\hat{z}^{2}}{2\sigma_{K}^{2}}}$
- $\int_{-\infty}^{-\hat{z}} f_{2}dz=\int_{\hat{z}}^{\infty} f_{2}dz=\frac{1}{2}\left( \sigma_{K}\sqrt{2\pi}\frac{K_{max}\cdot N}{cu_{2}\left( N+1 \right)^{2}}\left( \sigma_{K}^{2}-ncu_{2} \right)\cdot erfc\left( \frac{\hat{z}}{\sqrt{2\sigma_{K}^{2}}} \right) -2\hat{z}\sigma_{K}^{2}e^{\frac{-\hat{z}^{2}}{2\sigma_{K}^{2}}} \right)$

We now numerically search for optimal specialization $u_{2}$ such that $\frac{du_{2}}{dt}=0$. We do this by first fixing the parameters$K_{max}, n, c,$and $\sigma_{K}.$Then we calculate $\frac{du_{2}}{dt}$depending upon the table and calculations for a chosen$u_{2}$. The bisection method for root solving is then used over the interval $u_{2}=[0.001,10,000]$ (or for multiple intervals within that range to determine multiple points of optimization) to arrive at optimal specialization.

### **Calculating Disruptive Selection on Niche Position**

To calculate disruptive vs stabilizing selection on the population, we use the same process as was done with $\frac{du_{2}}{dt}$ only with $\frac{\partial^{2}G}{\partial v_{1}^{2}}$. Caluculating $\frac{\partial^{2}G}{\partial v_{1}^{2}}$, we get

$$\frac{\partial^{2}\hat{\alpha}(z_{j},\vec{v},\mathbf{u},\vec{N})}{\partial v_{1}^{2}}=\left\{ \begin{matrix} \frac{\partial^{2}\alpha(z_{j},\vec{v})}{\partial v_{1}^{2}} & ,\alpha_{T}(z_{j},\vec{v},\mathbf{u},\vec{N})<R(z_{j}) \\ \left( \frac{\partial^{2}\alpha\left( z_{j},\vec{v} \right)}{\partial v_{1}^{2}}-\frac{2\left( \frac{\partial\alpha\left( z_{j},\vec{v} \right)}{\partial v_{1}} \right)^{2}}{\alpha_{T}\left( z_{j},\vec{v},\mathbf{u},\vec{N} \right)} \right)\frac{\sum_{i=1}^{n_{s}} \alpha(z_{j},\vec{u_{i}})N_{i}}{{\alpha_{T}\left( z_{j},\vec{v},\mathbf{u},\vec{N} \right)}^{2}}\cdot R(z_{j}) & ,\alpha_{T}(z_{j},\vec{v},\mathbf{u},\vec{N})>R(z_{j}) \end{matrix} \right.$$

(4)

with $\frac{\partial\alpha(z_{j},\vec{v})}{\partial v_{1}}=\frac{2(z-v_{1})}{cv_{2}}\alpha(z_{j},\vec{v})$ and $\frac{\partial^{2}\alpha(z_{j},\vec{v})}{\partial v_{1}^{2}}=\frac{4(z_{j}-v_{1})^{2}-2cv_{2}}{(cv_{2})^{2}}\alpha(z_{j},\vec{v})$.

The table from earlier remains the same. Similar to before, let $g_{1}=\frac{\partial^{2}\alpha(z_{j},\vec{v})}{\partial v_{1}^{2}}\mid_{\vec{v}=\vec{u}}$ and $g_{2}=\left( \left( \sum_{i=1}^{n_{s}} \alpha\left( z_{j},\vec{u_{i}} \right)N_{i}\cdot\frac{\partial^{2}\alpha\left( z_{j},\vec{v} \right)}{\partial v_{1}^{2}}-2\left( \frac{\partial\alpha\left( z_{j},\vec{v} \right)}{\partial v_{1}} \right)^{2}+\alpha\left( z_{j},\vec{v} \right)\cdot\frac{\partial^{2}\alpha\left( z_{j},\vec{v} \right)}{\partial v_{1}^{2}} \right)\frac{\sum_{i=1}^{n_{s}} \alpha\left( z_{j},\vec{u_{i}} \right)N_{i}}{\alpha_{T}(z_{j},\vec{v},\mathbf{u},\vec{N})^{3}} \right)\mid_{\vec{v}=\vec{u}}\cdot R(z)$. If there are sufficient amounts of all resources, then $\frac{\partial^{2}G(\vec{v},\mathbf{u},\vec{N})}{\partial v_{1}^{2}}\mid_{\vec{v}=\vec{u}}=\int_{-\infty}^{\infty} g_{1}dz$. If there are insufficient amounts of all resources, then $\frac{\partial^{2}G(\vec{v},\mathbf{u},\vec{N})}{\partial v_{1}^{2}}\mid_{\vec{v}=\vec{u}}=\int_{-\infty}^{\infty} g_{2}dz$. If there are sufficient amounts of core resources and insufficient amounts of marginal resources, then $\frac{\partial^{2}G(\vec{v},\mathbf{u},\vec{N})}{\partial v_{1}^{2}}\mid_{\vec{v}=\vec{u}}=\int_{-\infty}^{-\hat{z}} g_{2}dz+\int_{-\hat{z}}^{\hat{z}} g_{1}dz+\int_{\hat{z}}^{\infty} g_{2}dz$. If there are insufficient amounts of core resources and sufficient amounts of marginal resources, then $\frac{\partial^{2}G(\vec{v},\mathbf{u},\vec{N})}{\partial v_{1}^{2}}\mid_{\vec{v}=\vec{u}}=\int_{-\infty}^{-\hat{z}} g_{1}dz+\int_{-\hat{z}}^{\hat{z}} g_{2}dz+\int_{\hat{z}}^{\infty} g_{1}dz$. Solving for each component reveals

- $\int_{-\infty}^{\infty} g_{1}dz=0$
- $\int_{-\infty}^{\infty} g_{2}dz=-2K_{max}N\sqrt{\frac{2\pi\sigma_{K}^{2}}{cu_{2}}}\frac{cu_{2}\left( N+1 \right)+2\sigma_{K}^{2}}{u_{2}^{n}\left( \left( N+1 \right)\sqrt{cu_{2}+{2\sigma}_{K}^{2}} \right)^{3}}$
- $\int_{-\hat{z}}^{\hat{z}} g_{1}dz=-\frac{4\hat{z}}{cu_{2}^{n+1}}e^{-\frac{\hat{z}^{2}}{cu_{2}}}$
- $\int_{-\infty}^{-\hat{z}} g_{1}dz=\int_{\hat{z}}^{\infty} g_{1}dz=\frac{2\hat{z}}{cu_{2}^{n+1}}e^{-\frac{\hat{z}^{2}}{cu_{2}}}$
- $\int_{-\hat{z}}^{\hat{z}} g_{2}dz =-\frac{2K_{max}N\sqrt{\frac{2\pi\sigma_{K}^{2}}{cu_{2}}}\left( cu_{2}\left( N+1 \right)+2\sigma_{K}^{2} \right)}{u_{2}^{n}\left( \left( N+1 \right)\sqrt{cu_{2}+{2\sigma}_{K}^{2}} \right)}\cdot erf\left( \frac{\hat{z}\left( \sqrt{2\sigma_{K}^{2}}+\sqrt{cu_{2}} \right)}{\sqrt{2\sigma_{K}^{2}cu_{2}}} \right)-\frac{8K_{max}N^{2}\hat{z}\sigma_{K}^{2}}{cu_{2}^{n+1}\left( cu_{2}+2\sigma_{K}^{2} \right)\left( N+1 \right)^{3}}e^{\frac{\hat{z}^{2}\left( 2\sigma_{K}^{2}+cu_{2} \right)}{2\sigma_{K}^{2}cu_{2}}}$
- $\int_{-\infty}^{-\hat{z}} g_{2}dz=\int_{\hat{z}}^{\infty} g_{2}dz=-\frac{K_{max}N\sqrt{\frac{2\pi\sigma_{K}^{2}}{cu_{2}}}\left( cu_{2}\left( N+1 \right)+2\sigma_{K}^{2} \right)}{u_{2}^{n}\left( \left( N+1 \right)\sqrt{cu_{2}+{2\sigma}_{K}^{2}} \right)}\cdot erfc\left( \frac{\hat{z}\left( \sqrt{2\sigma_{K}^{2}}+\sqrt{cu_{2}} \right)}{\sqrt{2\sigma_{K}^{2}cu_{2}}} \right)-\frac{4K_{max}N^{2}\hat{z}\sigma_{K}^{2}}{cu_{2}^{n+1}\left( cu_{2}+2\sigma_{K}^{2} \right)\left( N+1 \right)^{3}}e^{\frac{\hat{z}^{2}\left( 2\sigma_{K}^{2}+cu_{2} \right)}{2\sigma_{K}^{2}cu_{2}}}$

We now substitute all relevant parameters and optimal $u_{2}$ into the equations to calculate $\frac{\partial^{2}G(\vec{v},\mathbf{u},\vec{N})}{\partial v_{1}^{2}}\mid_{\vec{v}=\vec{u}}$ and determine if selection is disruptive $\left( \frac{\partial^{2}G\left( \vec{v},\mathbf{u},\vec{N} \right)}{\partial v_{1}^{2}}\mid_{\vec{v}=\vec{u}}>0 \right)$ or stabilizing $\left( \frac{\partial^{2}G\left( \vec{v},\mathbf{u},\vec{N} \right)}{\partial v_{1}^{2}}\mid_{\vec{v}=\vec{u}}<0 \right)$.
